## Supplementary material for "Sperm motility in mice with oligo-astheno-teratozoospermia restored by *in vivo* injection and electroporation of naked mRNA": video 6

### Shared first authorship.

* To whom correspondence should be addressed: Jessica Escoffier, Team "Genetics, Epigenetics and Therapies of Infertility", Institute for Advanced Biosciences (IAB), INSERM 1209, CNRS UMR 5309 University Grenoble Alpes, Grenoble, FRANCE.
Contact:mail to

**Key words:** Sperm cells, infertility, protein therapy, mRNA, EEV, *In Vivo* Microinjection and Electroporation, *in vivo* imaging, Whole Testis Optical clearing, lightsheet microscopy.

#### Introduction

Worldwide, 10-15 % of couples (or 70 million) face infertility [1]. Infertility is thus a major public health issue presenting significant medical, scientific and economic challenges (a multibillion € annual market)[2]. A significant proportion of infertilities is due to altered gametogenesis, where the sperm and eggs produced are incompatible with fertilization and/or embryonic development [3]. Approximately 40 % of cases of infertilities involve a male factor, either exclusively, or associated with a female deficiency [4].

For GFP expression analysis, the same program was used, except that the annealing temperature was set to 62°C instead of 60°C.

Relative gene expression was calculated using the 2^–ΔCT method.

| | | Primer sequences (5'-3') | Final concentration (µM) |
|---|---|---|---|
| Genotyping | | | |
| *Armc2*_WT | Forward | GGCCCGAGCACGCTTCTA | 0.4 |
| | Reverse | TTCATGTAAGAACTATCCAGGACCA | 0.4 |
| *Armc2*_KO | Forward | TGGGACGCAGCCCTGTAA | 0.4 |
| | Reverse | AACCCAAAGCTCCAGCATCTC | 0.4 |
| Expression | | | |
| *Gfp* | Forward | ACGACTTCTTCAAGTCCGCC | 0.08 |
| | Reverse | TCTTGTAGTTGCCGTCGTCC | 0.08 |
| *Gadph* Human | Forward | TCTCTGCTCCTCCTGTTCGA | 0.33 |
| | Reverse | TTCCCGTTCTCAGCCTTGAC | 0.33 |
| *Gadph* mice | Forward | | |
| | Reverse | | |

Table 1. Primer sequences used for mouse genotyping and *Gfp* expression analysis in HEK cells and mouse seminiferous tubules.
