## Supplementary material for "Sperm motility in mice with oligo-astheno-teratozoospermia restored by *in vivo* injection and electroporation of naked mRNA": manuscript revised version

**Tissue collection and histological analysis**

For histological analysis, treated and control B6D2 testes were fixed by immersion in 4 % paraformaldehyde (PFA) for 14 h. They were then embedded in paraffin before cutting into 5 µm sections using a microtome (Leica biosystems, [Wetzlar, Germany](https://www.google.com/search?sa=X&sca_esv=3f65f6f3feeae64f&sca_upv=1&rlz=1C1ONGR_frFR951FR951&biw=2133&bih=1050&sxsrf=ADLYWIKgVywKit9O1h7psWXHlYlydcIxqw:1726221119932&q=Wetzlar&si=ACC90nzx_D3_zUKRnpAjmO0UBLNxnt7EyN4YYdru6U3bxLI-L7mj9B_tpdX-9B7T0fgFfRfoGABXxrV7sVwTBL-3wH1k7Nn7aS6JnyFax1O_8kFuB_ZjQbZteISF3i9p3PFVCTBm4LWKjki50TxUIyEzRpNxXb9SMMkEh0Nda2EZYewW_aiLo2apaSOOUgyGQ72peAMgX-_h&ved=2ahUKEwjo0qDl0r-IAxV7SaQEHQRjMNYQmxMoAHoECEAQAg)). After deparaffination, the sections were stained with hematoxylin and eosin. Stained sections were digitized at 20x magnification using an Axioscan Z1-slide scanner (Zeiss, Jna, Germany). Spermatogenesis was assessed by measuring the area of seminiferous tubules and the cross sections of round tubules (μm^2^) (n > 35 tubules per testis section; n=5 testis sections per condition). Statistical significance of differences was determined using a Student’s t-test.

ICSI procedures

ICSI was performed according to the method described by [Yoshida and Perry (2007)](javascript:;)[33].For microinjection, *Armc2*^-/-^ sperm or *Armc2*^-/-^ -rescued motile sperm heads were separated from the tail by applying multiple piezo pulses (PiezoXpert^®^, Eppendorf, Montesson, France). Sperm heads were introduced into the ooplasm using micromanipulators (Micromanipulator InjectMan^®^, Eppendorf, Montesson, France) mounted on an inverted Nikon TMD microscope (Nikon, [Minato-ku, Tokyo, Japan](https://www.google.com/search?sca_esv=ea85460ec5208a83&sca_upv=1&rlz=1C1ONGR_frFR951FR951&sxsrf=ADLYWII-5o3Sc44yEdDdJVRJA6O9dUD6dg:1726227539583&q=Minato-ku&si=ACC90nzx_D3_zUKRnpAjmO0UBLNxnt7EyN4YYdru6U3bxLI-L6ybrC4rO_YmtM9g_AbT4NZSpdqFnXJHWVc_N5UqI7V3eP5dAN810TQ-iRu07KGLnAj8oq8kgmGMlIXgAa0miOuWrFPfQvhguoZ-IT5pXE5JdIZ3XGSw9eID0fcIyX_KJ6-2N7FOHE1eGzW-JzE2bY4rifQU&sa=X&ved=2ahUKEwiboLHa6r-IAxVXVKQEHWKZF3gQmxMoAHoECEEQAg)). Eggs that survived the ICSI procedure were incubated in KSOM medium at 37 °C under in an atmosphere of 5 % CO_2_. Pronucleus formation was checked at 6 h after ICSI, and outcomes were scored up to the blastocyst stage.
